# A persistent giant algal virus, with a unique morphology, encodes an unprecedented number of genes involved in energy metabolism

**DOI:** 10.1101/2020.07.30.228163

**Authors:** Romain Blanc-Mathieu, Håkon Dahle, Antje Hofgaard, David Brandt, Hiroki Ban, Jörn Kalinowski, Hiroyuki Ogata, Ruth-Anne Sandaa

## Abstract

Viruses have long been viewed as entities possessing extremely limited metabolic capacities. Over the last decade, however, this view has been challenged, as metabolic genes have been identified in viruses possessing large genomes and virions—the synthesis of which is energetically demanding. Here, we unveil peculiar phenotypic and genomic features of *Prymnesium kappa* virus RF01 (PkV RF01), a giant virus of the *Mimiviridae* family. We found that this virus encodes an unprecedented number of proteins involved in energy metabolism, such as all four succinate dehydrogenase (SDH) subunits (A–D) as well as key enzymes in the *β*-oxidation pathway. The *SDHA* gene was transcribed upon infection, indicating that the viral SDH is actively used by the virus— potentially to modulate its host’s energy metabolism. We detected orthologous *SDHA* and *SDHB* genes in numerous genome fragments from uncultivated marine *Mimiviridae* viruses, which suggests that the viral SDH is widespread in oceans. PkV RF01 was less virulent compared with other cultured prymnesioviruses, a phenomenon possibly linked to the metabolic capacity of this virus and suggestive of relatively long co-evolution with its hosts. It also has a unique morphology, compared to other characterized viruses in the *Mimiviridae* family. Finally, we found that PkV RF01 is the only alga-infecting *Mimiviridae* virus encoding two aminoacyl-tRNA synthetases and enzymes corresponding to an entire base-excision repair pathway, as seen in heterotroph-infecting *Mimiviridae*. These *Mimiviridae* encoded-enzymes were found to be monophyletic and branching at the root of the eukaryotic tree of life. This placement suggests that the last common ancestor of *Mimiviridae* was endowed with a large, complex genome prior to the divergence of known extant eukaryotes.

**Importance:** Viruses on Earth are tremendously diverse in terms of morphology, functionality, and genomic composition. Over the last decade, the conceptual gap separating viruses and cellular life has tightened because of the detection of metabolic genes in viral genomes that express complex virus phenotypes upon infection. Here, we describe *Prymnesium kappa* virus RF01, a large alga-infecting virus with a unique morphology, an atypical infection profile, and an unprecedented number of genes involved in energy metabolism (such as the tricarboxylic (TCA) cycle and the *β*-oxidation pathway). Moreover, we show that the gene corresponding to one of these enzymes (the succinate dehydrogenase subunit A) is transcribed during infection and is widespread among marine viruses. This discovery provides evidence that a virus has the potential to actively regulate energy metabolism with its own gene.

## Introduction

In their essay “Varieties of Living Things: Life at the Intersection of Lineage and Metabolism,” Dupré and O’Malley proposed to address Schrödinger’s question “What is Life?” by “*describing a spectrum of biological entities that illustrates why no sharp dividing line between living and non-living things is likely to be useful*” (1). Microbiologists have contributed considerably to this descriptive effort, both by reporting the existence of viruses endowed with genes coding for functions once thought to be exclusive to cellular life and by concomitantly proposing that actively infecting viruses are a “living form” (2–4). Genes encoding elements for photosynthesis (5, 6), carbon metabolism (7), and nitrogen- (8) and sulfur-cycling (9) have been found in bacterial viruses, where they are used to maintain or augment cellular processes during infection and to redirect energy and resources towards viral production (8, 10, 11). Genes for protein synthesis, including translation initiation, elongation, and termination, and a range of aminoacyl-tRNA synthetases have been found in *Mimiviridae*, a group of giant viruses infecting single-celled eukaryotes (12–14). *Mimiviridae* and other large DNA viruses, including some bacterial viruses, also have tRNA genes (15, 16). Ribosomal proteins have recently been reported in viral genomes derived from metagenomes (17). Genes involved in other metabolic processes, such as fermentation (18), glycosylation (19), photosynthesis (20), and rhodopsin (21), are encoded in *Mimiviridae* and other related large eukaryotic DNA viruses. Metabolic genes are frequently observed within virus genomes (20, 22, 23); although they represent a tiny fraction of the viral gene pool, these genes have the potential to dramatically modify the phenotype of an actively infected cell and alter the ecological role of the host (7, 24, 25). The infected host in this state has been referred to as a virocell (2). One might expect that the interplay between viral genes and host genes in virocells would become increasingly fine-tuned and complex during prolonged virus–host co-evolution, which also typically leads to lower virulence. Much of the complexity of virocells may still be undetected, as most *Mimiviridae* isolated with their natural host (mostly algae) are highly virulent, with several involved in rapid algal bloom termination events (26).

Viruses of the *Mimiviridae* family are known to infect heterotrophic and autotrophic microbial eukaryotes. This divide is also reflected in the phylogeny of these viruses, some of which are classified into two proposed sub-families: “Megavirinae” and “Mesomimivirinae” (27). The former contains viruses with genomes larger than 1 Mbp, all isolated from Amoebozoa, while the latter includes viruses with smaller genomes isolated from haptophyte algae of class Prymnesiophyceae. Several *Mimiviridae* members outside these two groups have been characterized to some extent as well, namely, viruses isolated from heterotrophs (*Cafeteria roenbergensis* virus, CroV; *Bodo saltans* virus, BsV; Choano virus), autotrophs (*Aureococcus anophagefferens* virus, AaV; Tetraselmis virus 1, TetV; *Pyramimonas orientalis* virus, PoV; *Prymnesium kappa* virus RF01, PkV RF01), a metazoan (Namao virus), and metagenomes (Klosneuviruses). The Mesomimivirinae sub-family includes viruses infecting bloom-forming hosts, such as *Phaeocystis pouchetii, Phaeocystis globosa*, and *Prymnesium parvum* (PpV, PgV Group I, and PpDVAV, respectively) (28–30); it also includes several viruses infecting *Haptolina ericina* and *Prymnesium kappa*, which normally do not form massive blooms but are present at low densities in seawater year round (31). In marine environments, viruses infecting low-density and non-bloom-forming algae may be the most common virus–host systems—that is, low-density hosts (non-blooming) and viruses that appear to have co-evolved in response to host growth strategy. Thus far, the only known representatives of such viruses are *Prymnesium kappa* viruses RF01 (PkV RF01) and RF02 (PkV RF02), *Haptolina ericina* virus RF02 (HeV RF02), and *Chrysochromulina ericina* virus (CeV 01B, infecting *Haptolina ericina*) (32, 33). Together with PgV, all of these viruses, except for PkV RF01, belong to the subfamily Mesomimivirinae on the basis of their monophyletic relationship and, in the case of PgV and CeV, a shared genomic similarity (27). In contrast, phylogenetic analysis of two partially sequenced marker genes has placed PkV RF01 deep inside the *Mimiviridae* clade, and characterization of its life cycle has revealed an atypical infection profile (33). Here, we report new phenotypic features as well as new viral functions inferred from analysis of the genome sequence of PkV RF01. We found that this virus has a unique morphology, is less virulent than most other alga-infecting viruses and possesses an unprecedented number of energy-generating genes. We uncovered clues suggesting that members of *Mimiviridae* that potentially modulate the metabolism of their hosts are widespread in the ocean. Our findings of peculiar genomic features in a persistent virus provide new insights on virus–host coevolution and may stimulate further advances in modeling the history of their interaction.

## Results and Discussion

### PkV RF01 has an atypical morphology

The icosahedral PkV RF01 particle is approximately 400 nm in diameter (Fig. 1A-B). Beneath the capsid, several convoluted inner membranes fill approximately 66% of the interior. Treatment of chloroform can be used to identify possible functions of lipid membranes, as it acts to remove lipid molecules that might be essential for successful infection (34). Some algal viruses in the NCLDV group are sensitive to chloroform (30, 35, 36) with the suggestions that lipid containing inner or outer membranes are involved in the infection process (35, 37). In our experiment, chloroform treatment of PkV RF01 drastically reduced the infectivity of the virus. (Fig. 1C). As no outer membrane was detected by cryo-electron tomography, the sensitivity to chloroform might be linked to lipid components in either the capsid or the inner convoluted membranes. Internal lipid-containing membranes have been detected in several icosahedral-shaped double-stranded DNA viruses, including algal viruses belonging to families *Phycodnaviridae* and *Mimiviridae*, mimiviruses, and various bacteriophages (38–43). In all of these viruses, the inner membranes are suggested to play a role in the release of the viral nucleoprotein core or genome by fusing with the host plasma membrane (40, 42, 43). Inner membranes in currently described NCLDVs more or less adopt the icosahedral morphology defined by the outer layer of capsomers (44, 45). We detected several convoluted inner membranes in PkV RF01 that do not follow the structure of the capsid. To our knowledge, this structural inconsistency has not been previously detected in any double-stranded DNA viruses, which calls for further investigation to understand the assembly process of PkV RF01 and how it enters its host. Another striking feature of the PkV RF01 virion is an internal rod-shaped core (ca. 55 nm in diameter), which is filled with dense material and positioned in the center of the virus particle. Similar features have been observed in TEM images of large virus-like particles (VLPs) (300–700 nm) occurring in waste vacuoles of phaeodarian radiolarians collected from different oceans (46) and in zoospores of the green alga *Chlorococcus minutum* (47). To our knowledge, however, these features have not been described in isolated viruses thus far.

**FIG 1.**
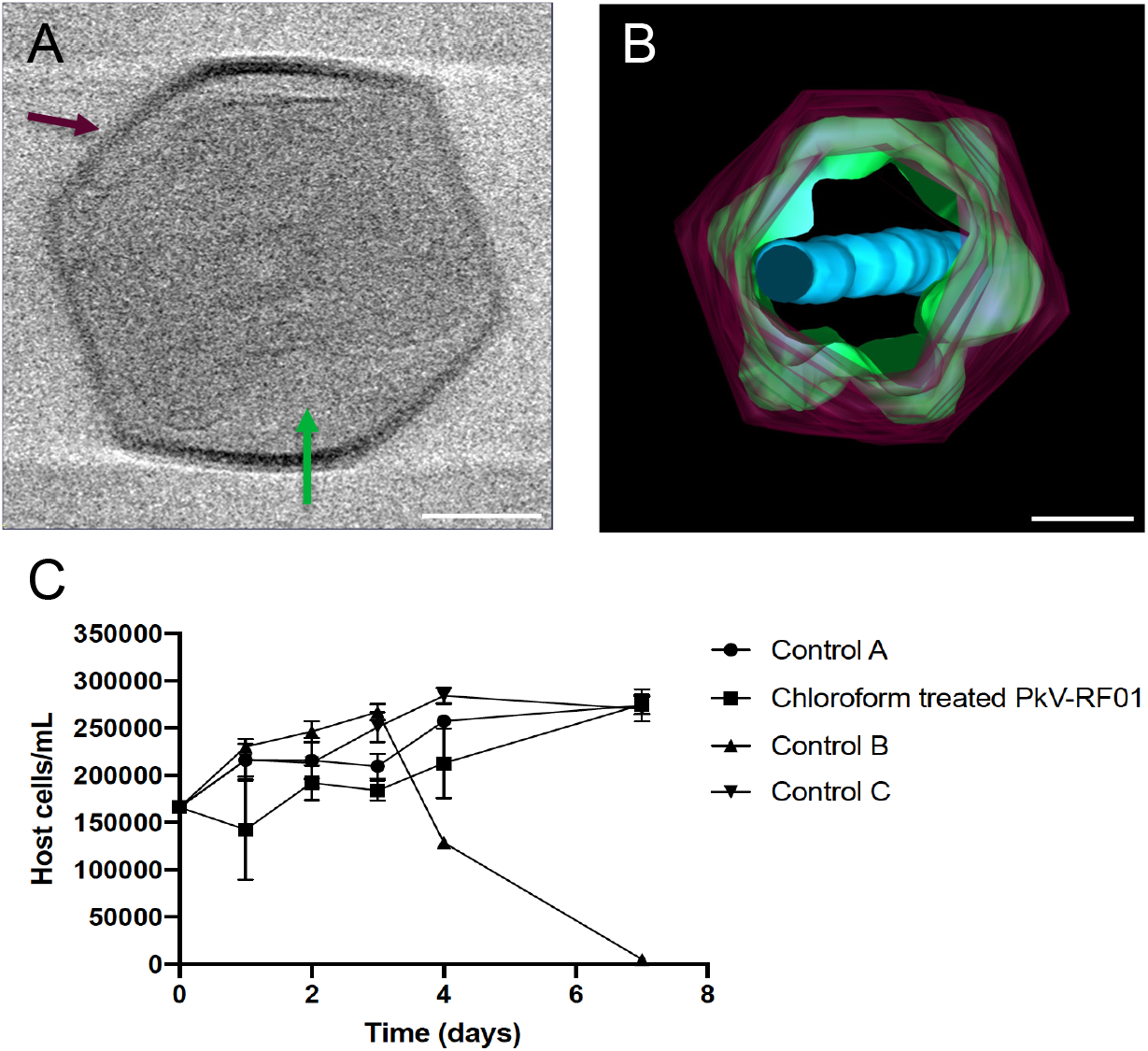
PkV RF01 morphology and reduced viral infectivity under chloroform treatment. (A) Screen shot of a cryo-electron tomogram of a PkV RF01 virion. (B) Composite image of 61 cryo-electron tomograms (−60 to 60°, imaged every 2°). Purple, capsid; green, inner membrane consisting of multiple irregular, convoluted membranes; blue, internal rod-shaped core filled with dense material. The full set of records is available on GitHub (see Data availability section). Scale bar, 100 nm. (C) Reduction of PkV RF01 infectivity with chloroform. Experiments were set up in triplicate, and host cells were counted by flow cytometry. Chloroform-treated PkV RF01 was added to exponentially growing He UiO028 cells in a 1:10 volume ratio. Controls were He UiO028 cells incubated with chloroform-treated medium (Control A), untreated PkV RF01 (Control B), and untreated medium (Control C). SDs are indicated with error bars.

### PkV RF01 has an atypical infection strategy

Only 2% of the total PkV RF01 viral particles produced during infection of *Haptolina ericina* UiO028 (He UiO028) were infectious (able to produce progeny) (Table 1). This infectivity was much lower than that of the other two prymnesioviruses, HeV RF02 and PkV RF02, which produced 13% and 44% of infectious progeny respectively (Table 1). The portion of infectious particles of PkV RF01 is low also when compared to other algal viruses (48, 49). In addition, the latent period of PkV RF01 was previously reported to be longer (ca. 24–32 h, (33)) in comparison with other prymnesioviruses (28, 29, 32, 33) and it has been demonstrated that PkV RF01 is also able to infect multi-species (33), that is another unusual trait among algal viruses (26).

**TABLE 1.** Infection parameters of *Prymnesium kappa* viruses RF01 and RF02 and *Haptolina ericina* virus RF02.

| Viral species and hosts | Infectious progeny/mL (MPN) | Host cells/mL (FCM)<sup>a</sup> | Total VLP/mL (FCM) | Burst size (VLP)<sup>b</sup> | Infectivity (%)<sup>c</sup> | Infectious particles in a burst<sup>d</sup> |
| --- | --- | --- | --- | --- | --- | --- |
| PkV RF01 (He UiO028) | 2.9x10<sup>6</sup> (± 0.2) | 4.9x10<sup>5</sup> | 1.8x10<sup>8</sup> (±0.9) | 363 | 2 | 6 |
| PkV RF02 (Pk RCC3423) | 2.2x10<sup>8</sup> (± 0.2) | 4.6x10<sup>5</sup> | 5.0x10<sup>8</sup> (±0.1) | 1093 | 44 | 483 |
| HeV RF02 (He UiO028) | 5.8x10<sup>7</sup> (±0.2) | 4.9 x 10<sup>5</sup> | 4.4 x10<sup>8</sup> (±0.0) | 907 | 13 | 119 |

VLP, virus-like particle; MPN, most probable number; FCM, flow cytometry.
<sup>a</sup>Measurement performed in duplicates
<sup>b</sup>The number of viral particles released from each host cell, estimated from the total number of host cells pre-infection and the total number of VLPs produced during the infection cycle.
<sup>c</sup>Estimated as the percentage of infectious progeny of all VLPs produced during the infection cycle.
<sup>d</sup>Number of infectious particles released per host cell.

The hosts of PkV RF01, PkV RF02, and HeV RF02 all belong to order the Prymnesiales, whose members are normally present in low abundance but co-occur year round (*K*-strategists) (50). PkV RF01, PkV RF02, and HeV RF02 are less virulent, as shown in the present study, and have longer latent periods compared with viruses infecting bloom-forming haptophytes (*r*-strategists). Two of these viruses (PkV RF01 and HeV RF02) are also able to infect multi species (generalists) (33). Longer replication time and reduced virulence, as hosts becomes scarce, increases the chances of vertical transmission rather than horizontal transmission of a virus. As vertical parent-to-offspring transmission depends on host reproduction, it has been argued that such transmission should select for reduced virulence because the virus depend on host survival and reproduction for its transmission (51, 52). High virulence, on the other hand, may be supported by large, dense host populations, as e.g. algal blooms, because high host densities ensure successful horizontal transmission of viral progeny to new hosts (51, 53). Viruses infecting the recurrent bloom-forming haptophytes, *Phaeocystis pouchetii* virus (PpV), and *Phaeocystis globosa* virus (PgV), are indeed highly virulent with between 60%–100% of virus particles produced being infectious, resulting in rapid lysis of their hosts (48, 54). Broad host range might also increase the chance of transmission in an environment with low host abundances (*K*-strategists). Such strategy requires a tradeoff whereby the virus decreases its opportunity of transmission by evolving longer replication times, higher decay rates and reduced infectivity (discussed in (55, 56)). This fits well with our two multi-species infecting haptophyte viruses, PkV RF01 and HeV RF02, that have reduced proportions of infectious particles and longer replication times (33), relative to other haptophyte viruses with restricted host ranges (specialists) like e.g. the *Emiliania huxleyi* virus (EhV), PpV and PgV.

The balance between fitness traits, such as virulence, latent period and host range, and tradeoffs is the result of the adaptive evolution between viruses and their hosts, resulting in relationships spanning from acute to stable coexistence (persistence). In the ocean, persistent relationships—such as between PkV RF01 and its hosts—seem to be most common among viruses infecting unicellular algae; this has been demonstrated by several metabarcoding studies revealing the persistence of dominance of viral OTUs over several months (57, 58). The atypical infection strategy of PkV RF01 evokes a persistent nature, different than the vast majority of other so far characterized algal viruses.

### PkV RF01 has the largest genome among algal viruses

The genome of PkV RF01 was assembled as a linear DNA sequence of 1,421,182 bp. This size is more than twice that of the genome of TetV, which means that PkV RF01 has the largest reported genome of any virus infecting a photosynthetic organism (Fig. 2A). Evidence for the linear structure of this genome is the presence of ~5-kbp terminal inverted repeats. Despite being phylogenetically more closely related to alga-infecting *Mimiviridae*, the genome size of PkV RF01 is in the range of heterotroph-infecting *Mimiviridae*. The overall G+C content of PkV RF01 is 22.8%, which is low compared with other *Mimiviridae* (23%–41%). Similar to other *Mimiviridae*, the average G+C content of PkV RF01 in intergenic regions is relatively low, 17.8%. This lower G+C content may reflect an ongoing loss of G and C nucleotides, more prevalent in non-coding than coding regions because of weaker background selection in non-coding regions. The genome of PkV RF01 is predicted to contain 1,161 genes comprising 1,121 protein-coding DNA sequences (CDSs) and 40 tRNA genes corresponding to 13 amino acids. Most tRNA genes (30 out of 40) are clustered in three genomic regions that lack predicted CDSs, a feature also observed in other *Mimiviridae*. For example, all tRNAs of TetV (*n* = 10) and CroV (*n* = 22) are encoded consecutively on the same strand (18, 59). The average CDS length is 1,046 bp (minimum: 297; maximum: 1,493). Intergenic regions average 217 bp in length, with a cumulative sum of 244,005 bp, which corresponds to a gene density of 82.8%.

**FIG 2.**
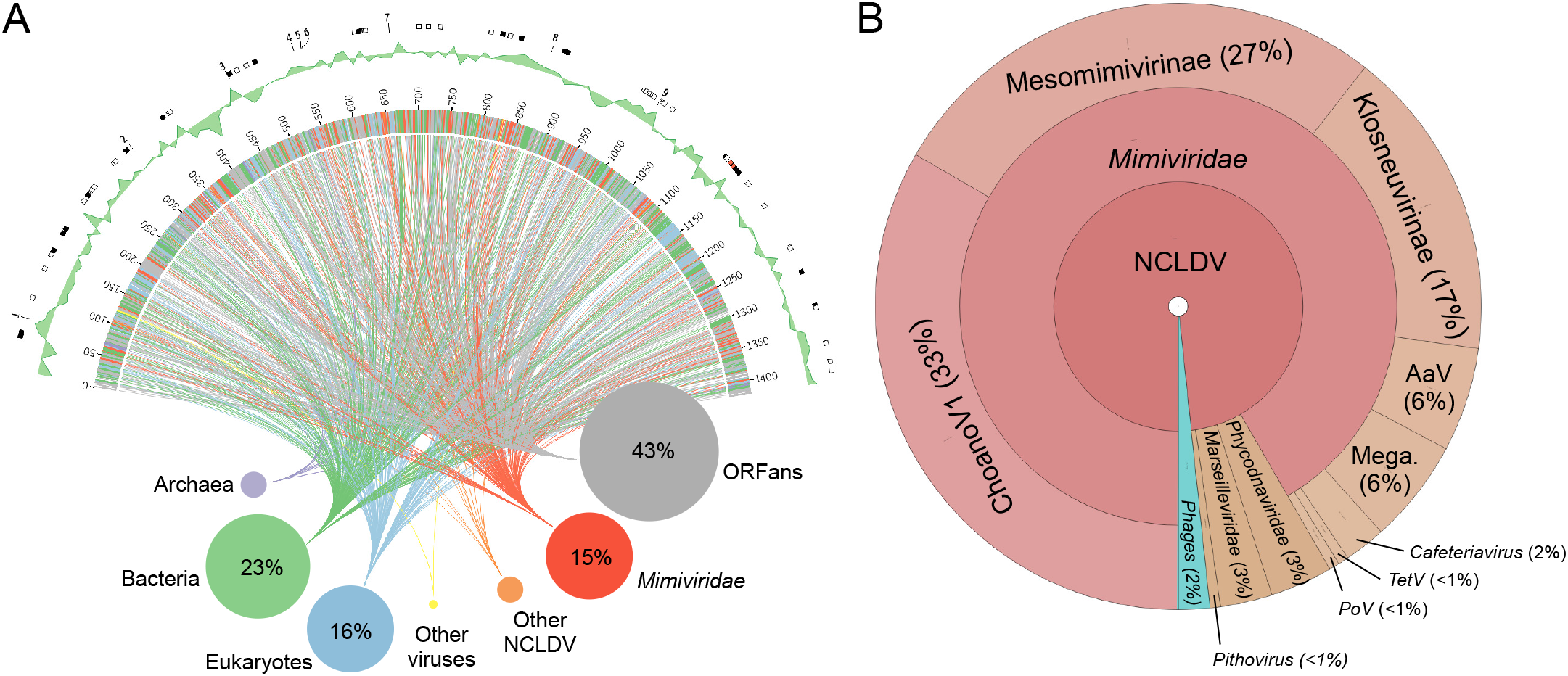
Structure and gene taxonomic composition of the PkV RF01 genome sequence. (A) Rhizome and genomic features of the PkV RF01 genome. As illustrated by the rhizome (inner part of the figure), ORFans comprise the largest set of PkV RF01 genes, and a substantial portion (15%) have their best BLAST hits (in UniRef90) against “*Mimiviridae*.” Colors indicate taxonomic origin. Intergenic regions are white. Percentage hits per taxonomic group higher than 5% of total genes are indicated. In the outermost ring, rectangles indicate the positions of glycosyltransferases (white), lipid-related enzymes (black), and succinate dehydrogenase genes (red), and the numbers correspond to *Mimiviridae* key enzymes (1 and 3: DNA-directed RNA polymerase II subunits 1 and 2, respectively; 2: DNA mismatch repair protein MutS7; 4: Packaging ATPase; 5: VLTF3, 6: Major capsid protein; 7: Eukaryotic translation initiation factor 4E; 8: Asparagine synthase; 9: DNA polymerase family B). The ring adjacent to the outermost ring shows GC skew over a 10-KB window. (B) Taxonomic breakdown of 180 genes with best hits to virus genes. Mega, Megavirinae; AaV, Aureococcus anophagefferens virus; TetV, Tetraselmis virus 1; PoV, Pyramimonas orientalis virus.

Of the 1,121 predicted CDSs, 641 (57%) exhibited sequence similarities (BLASTP *E*-value conservative cutoff of 1 × 10^-5^) to protein sequences in the UniRef90 database (Fig. 2A). Among them, 165 were most similar to *Mimiviridae*. Curiously, among the CDSs most similar to *Mimiviridae*, sixty were closest to ChoanoVirus which was isolated from choanoflagellates cultures, followed by Mesomimivirinae (*n* = 49) and Klosneuvirinae (*n* = 30) (Fig. 2B). Among the 181 closest homologs found in eukaryotic organisms 23 were haptophytes. A sequence-based homology search of corrected nanopore reads and scaffolds composing the initial assembly against *Lavidaviridae* proteomes (BLASTX; matrix: BLOSUM45, *E*-value < 1 × 10^-5^) yielded no significant alignments against any major or minor *Lavidaviridae* capsid proteins, which suggests that virophages were absent from the sample used for sequencing.

A previous analysis of PkV RF01 family-B DNA polymerase (PolB) and the major capsid protein (MCP) placed this virus in the family *Mimiviridae* (33). We also recently reported that the PkV RF01 genome has additional NCLDV core genes, such as A32-like virion packing ATPase (NCVOG0249) and RNApol (RNA pol subunit I [NCVOG0274] and subunit II [NCVOG0271]), and orthologous genes that are specific to *Mimiviridae*, namely, MutS7 (NCVOG2626) and asparagine synthase (AsnS, NCVOG0061) (60). Phylogenetic reconstruction using five NCLDV core genes confirmed the deep branching of PkV RF01 within the *Mimiviridae* family and suggested that PkV RF01, along with ChoanoV1, TetV and AaV, is more closely related to Mesomimivirinae than to Megavirinae (Fig. 3A). In support of this evolutionary relationship, PkV RF01 has an additional copy of the second largest RNA polymerase subunit gene (*rpb2*). This *rpb2* duplication is shared with all other *Mimiviridae* infecting algae, including Mesomimivirinae members, AaV (whose second copy is very short), and TetV and was previously proposed as a useful feature to discriminate between the two main clades (autotroph versus heterotroph-infecting viruses) within the *Mimiviridae* family (27). This additional *rpb2* copy is not found in other *Mimiviridae* to the exception of ChoanoV1 whose genome was derived from a single cell metagenome in choanoflagellates cultures. Phylogenetic analysis indicates that these two *rbp2* copies were present in the ancestor of alga-infecting *Mimiviridae* and ChoanoV1 (Fig. 3B). In agreement with the five NCLDV core genes phylogeny, it suggests that PkV RF01 and ChoanoV1, although evolutionarily distant, are more related with each other compared to any other *Mimiviridae*.

**FIG 3.**
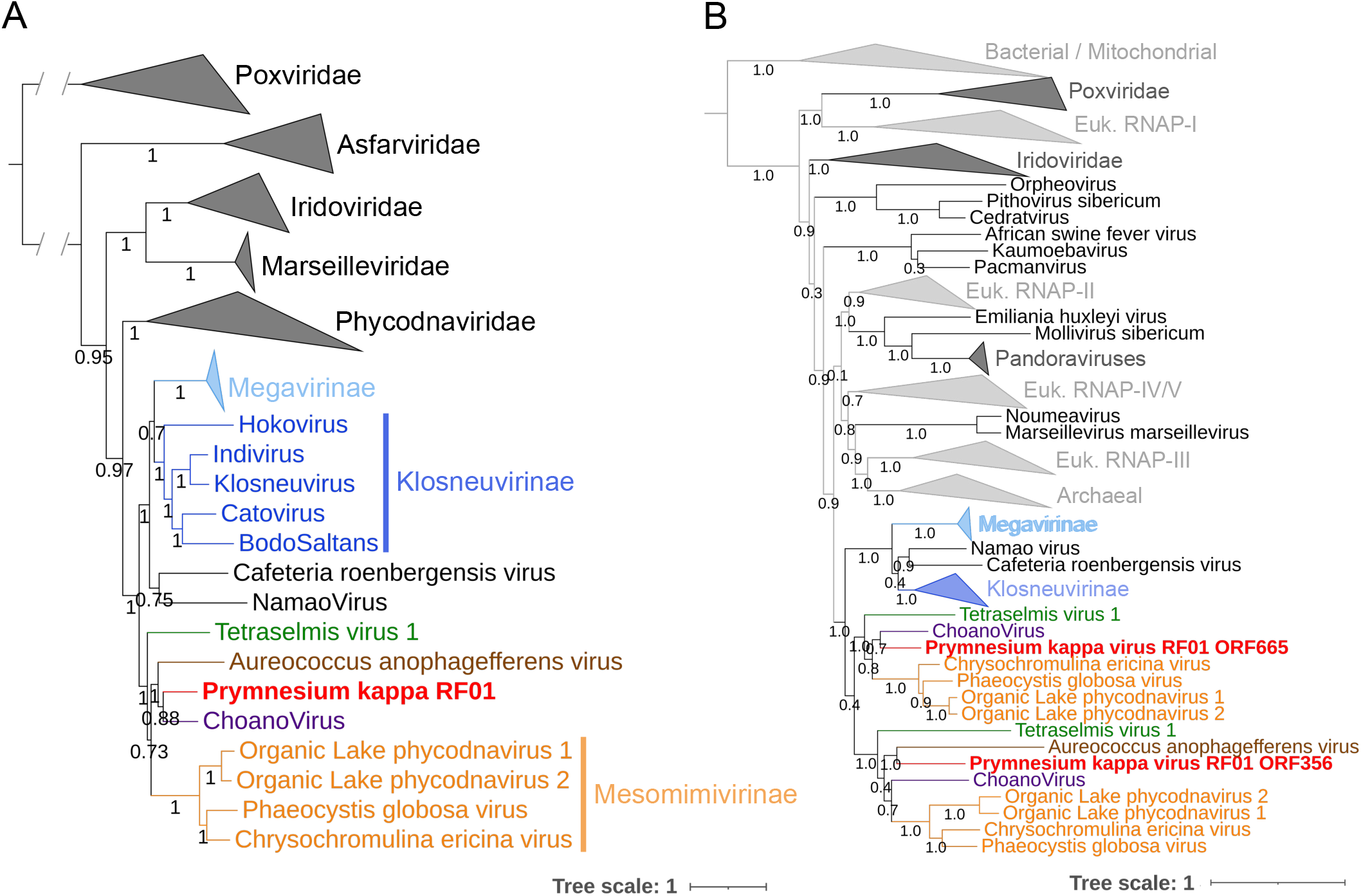
Phylogenetic evidence for PkV RF01 as a distant relative of “Mesomimivirinae.” (A) Bayesian phylogenetic tree of NCLDVs reconstructed from a concatenated alignment of five core nucleocytoplasmic virus orthologous genes. Values at branches are posterior probabilities support. The tree was rooted using *Poxviridae* as outgroup. The scale bar indicates substitutions per site. (B) Maximum likelihood phylogenetic tree of cellular and NCLDV DNA-directed RNA polymerase subunit beta (RPB2). Values at branches are Shimodaira-Hasegawa-like local support.

Out of 1,121 predicted protein-coding genes in the genome of PkV RF01, only about a third could be annotated with some functional description based on their sequence homology with characterized proteins. Such a small percentage is typical of divergent eukaryotic viruses detected for the first time. A total of 339 proteins (30%) showed significant sequence similarity with proteins in the Cluster of Orthologous Gene (COG) database (61) (Fig. 4). The distribution of COG functions associated with these hits was dominated by “Posttranslational modification, protein turnover, chaperones” (43 proteins) and “Cell wall/membrane/envelope biogenesis” (42 proteins), which is approximately two times more proteins than in other *Mimiviridae* members except for Tupanvirus. Among other well-represented categories, numbers of proteins in “Replication, recombination and repair” (36 proteins) and “Transcription” (23 proteins) were similar to those of other *Mimiviridae*, while the categories of “Translation, ribosomal structure and biogenesis” (25 proteins) and “Amino acid transport and metabolism” (20 proteins) were respectively in the same range or higher than those of heterotroph-infecting *Mimiviridae* (mimiviruses, BsV, and CroV). Interestingly, 24, 17, and 9 PkV RF01 proteins were respectively assigned to the categories of “Lipid transport and metabolism”, “Carbohydrates transport and metabolism,” and “Energy production and conservation,” all much higher compared with other *Mimiviridae* viruses (Fig. 5).

**FIG 4.**
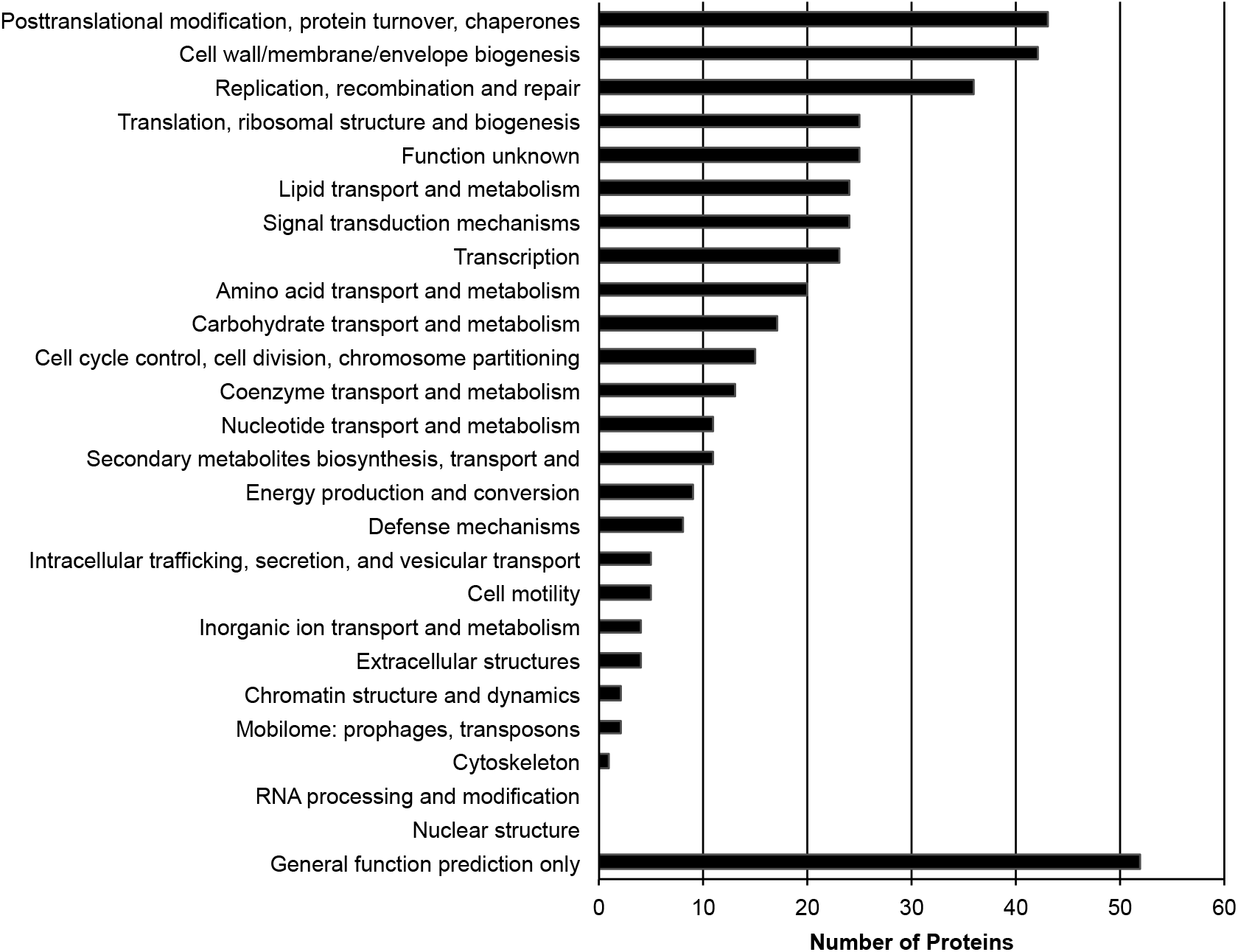
COG functional distribution of 339 proteins encoded by PkV RF01.

**FIG 5.**
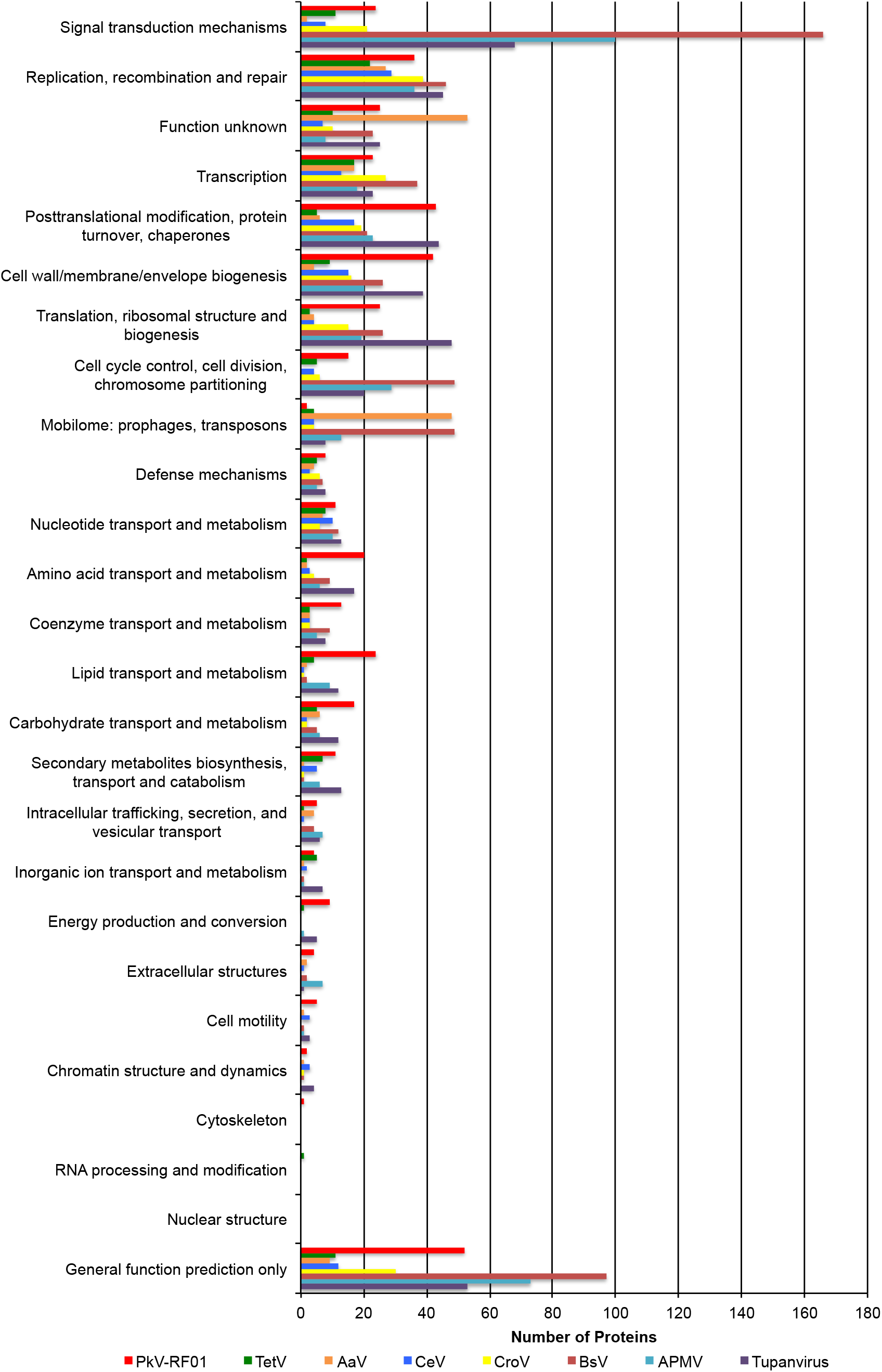
Comparative COG functional distribution among Mimiviridae members. COG sequences were automatically searched against the proteomes of each virus using BLASTP with an E-value of 1 × 10-5 as the significant similarity threshold.

Similar to other *Mimiviridae*, PkV RF01 encodes several genes involved in DNA repair, transcription, and translation. Notably, this virus has the full set of enzymes required for the base excision repair (BER) pathway, which is also the case for all *Mimiviridae* members except for those with smaller genomes (PgV, CeV, and AaV). PkV RF01 BER enzymes are closer (i.e., have a greater alignment score) to heterotroph-infecting *Mimiviridae* than to cellular homologs, thus suggesting that this pathway was present in the last common ancestor of *Mimiviridae*. According to a previous phylogenetic analysis, *Mimiviridae* BER enzymes are monophyletic with regard to *Mimiviridae* and have not recently been acquired from eukaryotes (62).

Unlike alga-infecting *Mimiviridae*, PkV RF01 encodes two amino-acyl tRNA synthetases (aaRS): an isoleucyl-tRNA synthetase (IleRS; ORF 480) and an asparaginyl-tRNA synthetase (AsnRS; ORF 764). Both of these synthetases are found in most lineages of heterotroph-infecting *Mimiviridae* (AsnRS is missing from CroV and BsV, and IleRS is missing from *Mimivirus* lineage A). Phylogenetic analyses of these two proteins revealed a deep branching of viral homologs, which formed a monophyletic clade well separated from cellular homologs (Fig. 6).

**FIG 6.**
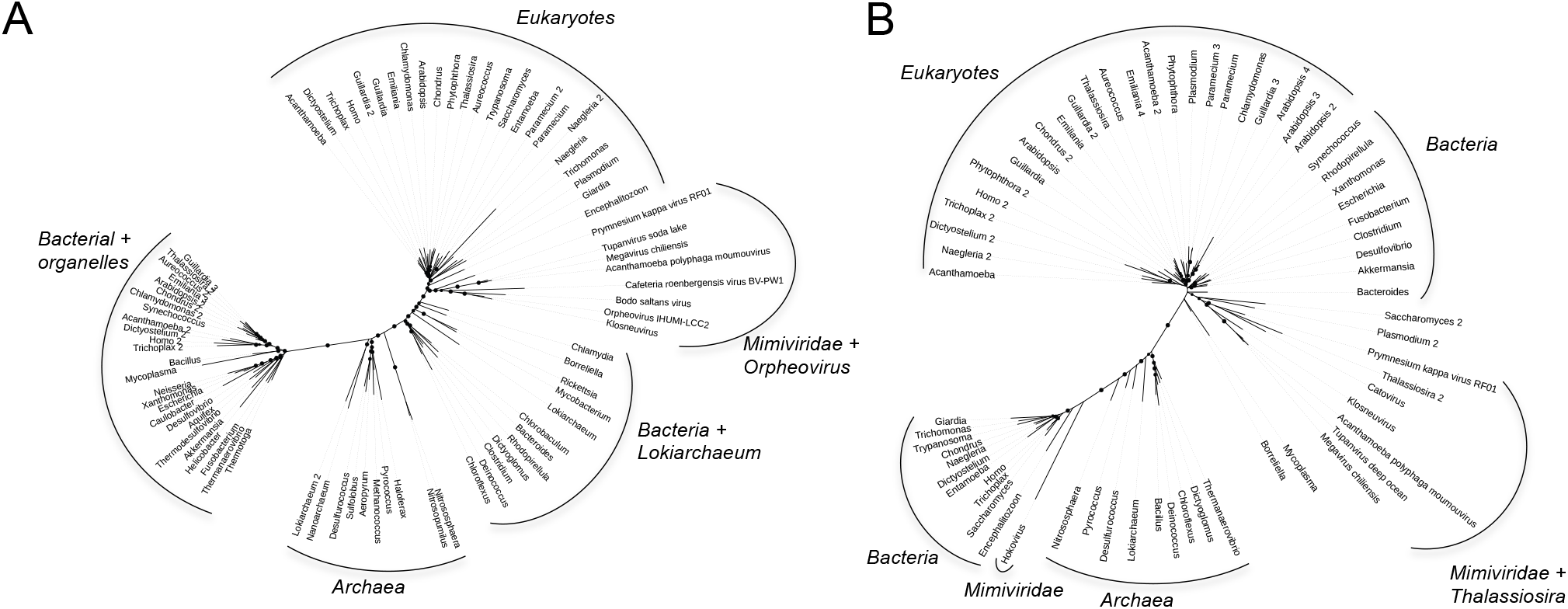
Bayesian phylogenetic trees of two viral amino-acyl tRNA synthetases and their cellular homologs. (A) Isoleucine tRNA synthases. (B) Aspartyl tRNA synthetases. Branches supported by posterior probability (PP) values >70% are indicated by circles whose diameters are proportional to the PP value.

### A viral-encoded succinate dehydrogenase and energy production genes

We found six predicted protein-coding genes (ORFs 893 to 900) related to energy production in an 8,026-bp region (Fig. 7A). Four ORFs (ORFs 893 and 898–900) were predicted to code for all four subunits (SDHA, D, C, and B) of a functional succinate dehydrogenase (SDH, or Electron Transport Chain Complex II) of the oxidative phosphorylation pathway (Fig. 7B). In eukaryotes, all four subunits of this enzyme are encoded in the nuclear genome. This enzyme acts in the mitochondrial respiratory chain and participates in both the TCA cycle and the respiratory electron transfer chain. In the TCA cycle, this succinate dehydrogenase oxidizes succinate to fumarate, while its activity in the inner mitochondrial membrane involves the reduction of a FAD cofactor followed by electron transfer through three Fe–S centers to ubiquinone (Fig. 7C).

**FIG 7.**
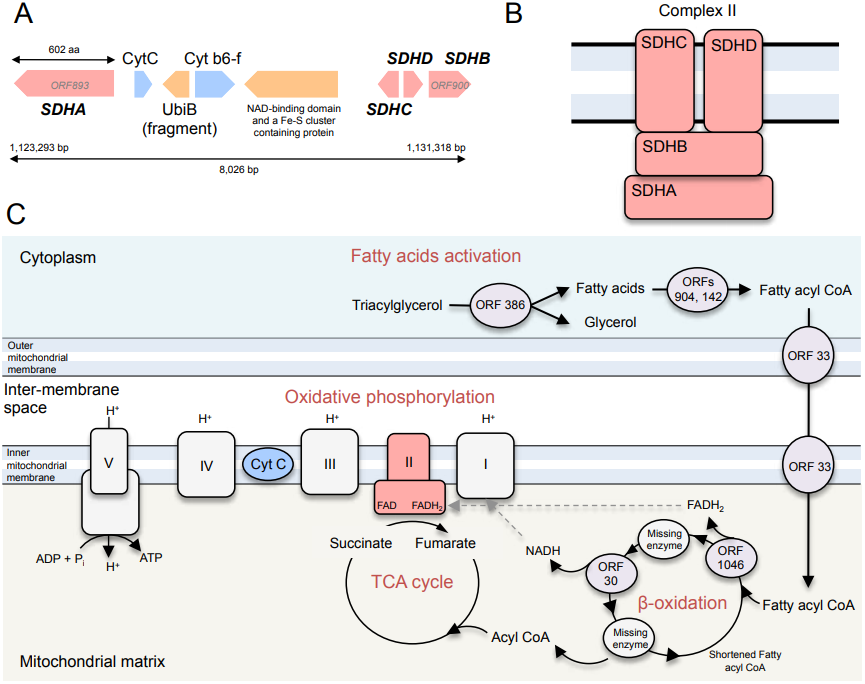
Genes in PkV RF01 predicted to encode enzymes of oxidative phosphorylation and β-oxidation pathways. (A) Gene organization in the succinate dehydrogenase-containing region. (B) Schematic representation of the canonical enzymatic complex II in the mitochondrial membrane. (C) Location of succinate dehydrogenase in the TCA cycle and electron transport chain as known in plants and a schematic reconstruction of the PkV RF01-encoded β-oxidation metabolic pathway.

*SDH* genes have recently been reported in viral genomes assembled from environmental samples for which functional experiments cannot be done (63). In a RT-PCR experiment using primers specific for the PkV RF01 gene for SDHA (hereafter, *vSDHA*), we detected transcripts of this gene in samples collected 24, 72, and 96 h post infection (Fig. 8). The *vSDHA* primers were tested on an uninfected culture to ensure that only the viral version of the *SDHA* gene was amplified (Fig. 9). The MCP gene of PkV RF01 was used both for protocol optimization and later as an internal positive control (Fig. 10). Although the transcription of the viral *SDHA* suggests that the viral SDH is functional, we can only speculate on the possible role of this enzyme during infection. One possibility is that the viral SDH sustains the carbohydrate metabolism of infected cells (i.e., virocells) to supply building blocks of viral particles such as amino acids and to support proper replication of this large virus. Another possibility is that PkV RF01 uses its SDH as a part of an arms race with its host to turn on the TCA cycle after the host had turned it off to counter viral replication, or more simply to boost the energy metabolism of the virocells to augment the fitness of the host and/or to maximize virus production efficiency.

**FIG 8.**
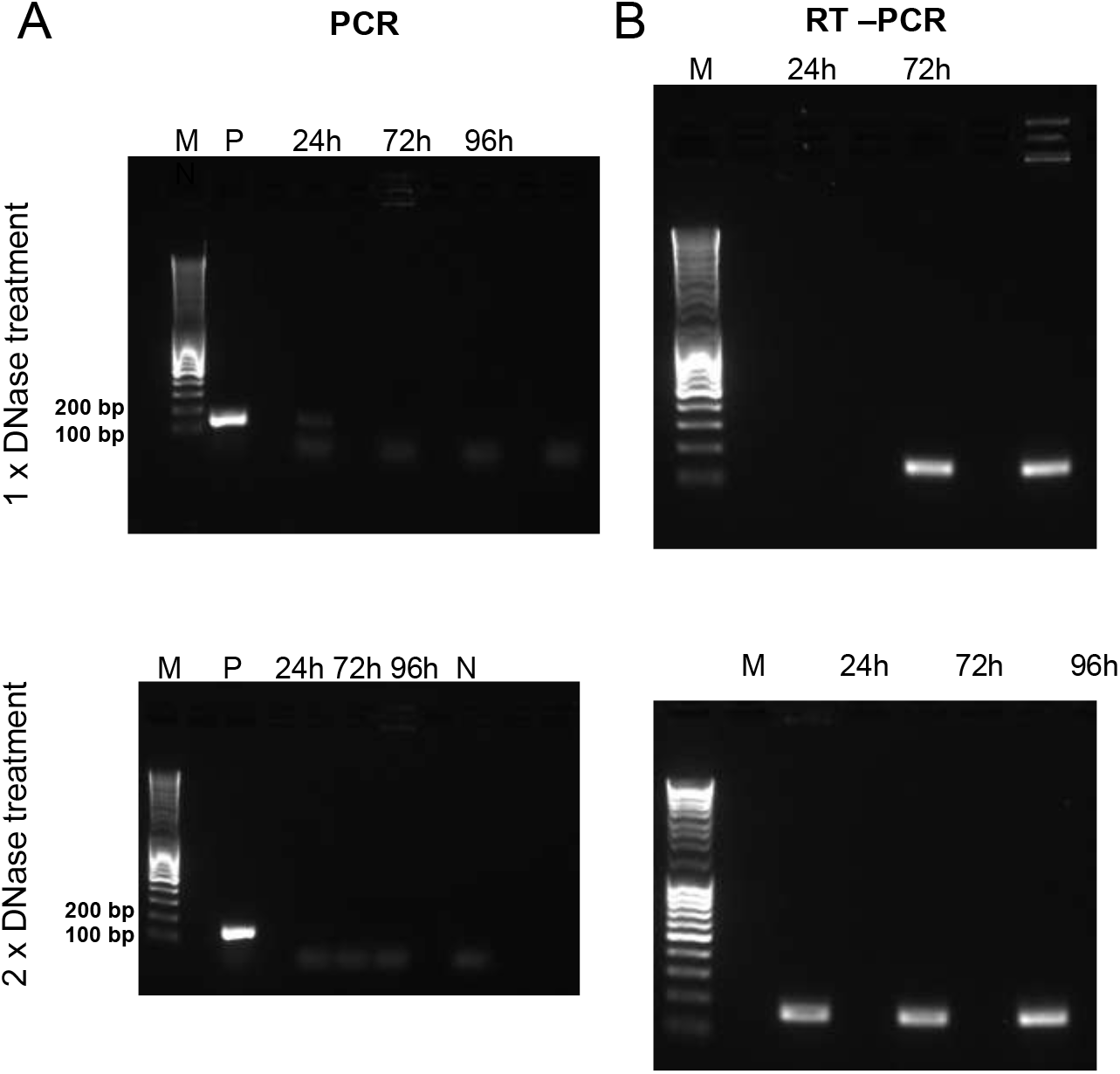
The viral SDHA gene is transcribed during infection. Gels of PCR and RT-PCR in combination with a TURBO DNA-free kit. Samples were taken 24, 72, and 96 h after infection. (A) PCR with *vSDHA*-specific primers was used to check for the presence of genomic DNA after RNA isolation treated with 1x and 2x DNAse, in the upper and lower panels respectively. P, positive control (PKV RF01 genomic DNA); N, negative control (sdH_2_0). (B) RT-PCR of RNA samples using *vSDHA*-specific primers. M, DNA marker (MassRuler DNA Ladder Mix, Thermo Fisher, 80 to 10,000 bp).

**FIG 9.**
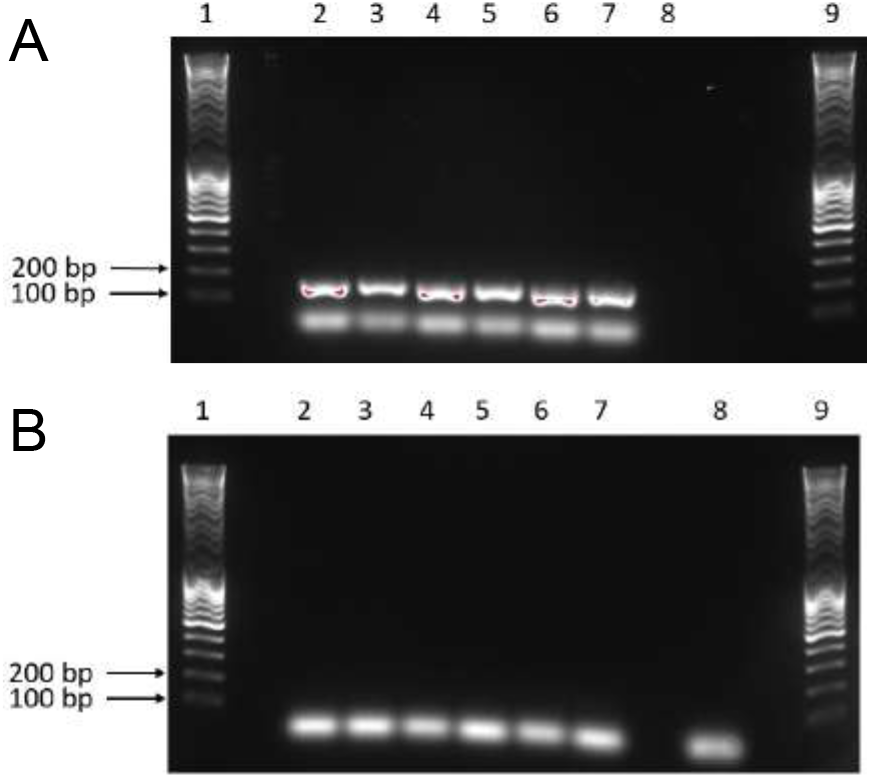
PCR optimization and confirmation of the SDHA gene in the PkV RF01 genome. (A–B) Results of PCR with SDHA primers using genomic PkV RF01 DNA (A) and genomic He UiO028 DNA (B) as templates. Lanes 1 and 9, DNA ladder; 2–7, optimization of the PCR annealing temperature from 55°C (2) to 60°C (7); 8, negative control (sdH2O).

**FIG 10.**
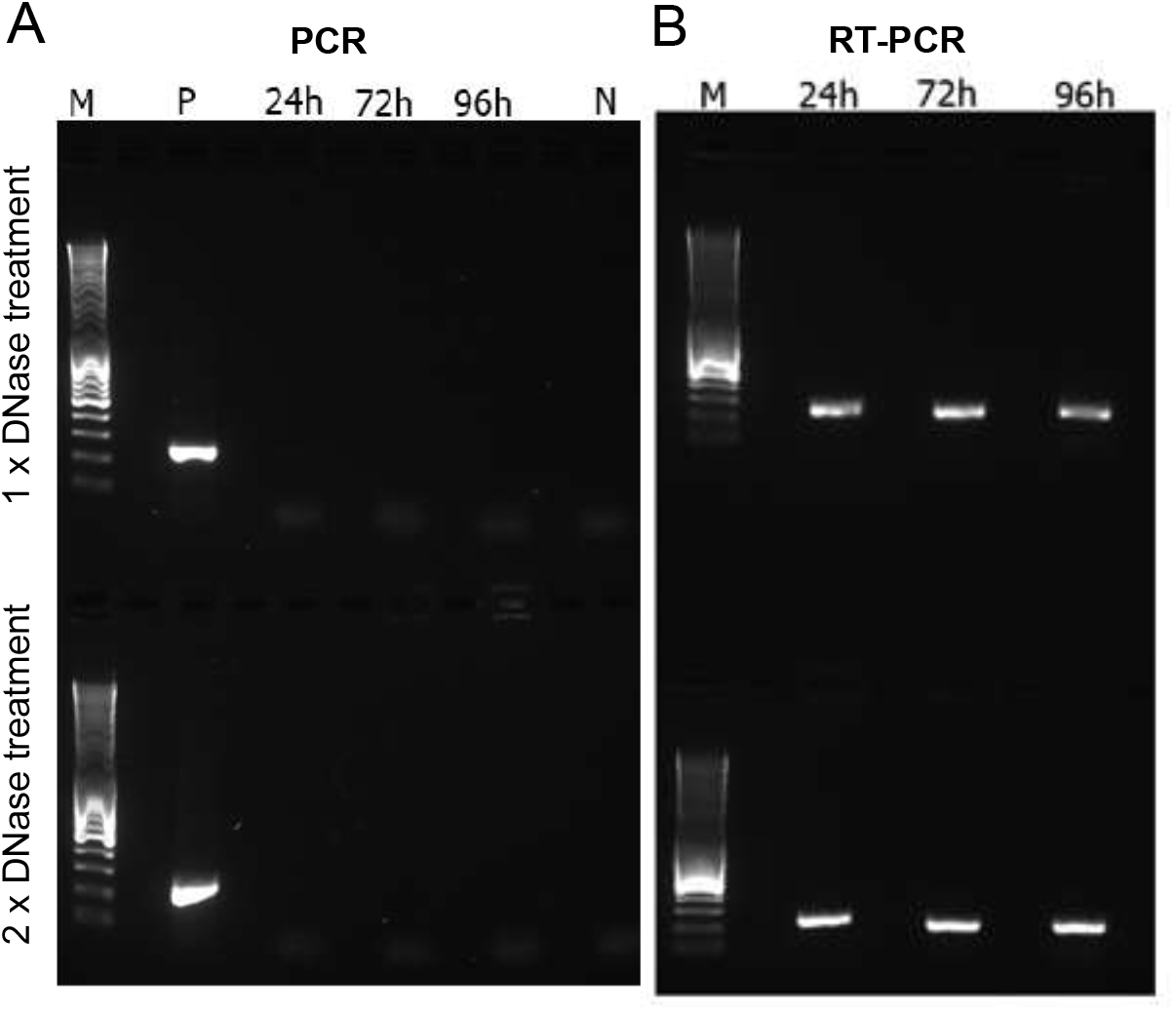
PCR and RT-PCR optimization using an internal control gene (mcp). PCR and RT-PCR were carried out after removal of genomic DNA using a TURBO DNA-free kit. Samples were taken 24, 72, and 96 h after infection. Two different protocols, both provided in the TURBO DNA-free kit manual, were used to optimize the reactions. (A) PCR check for the presence of genomic DNA after RNA isolation treated with 1x and 2x DNAse, in the upper and lower panels respectively. P, positive control (PkV RF01 genomic DNA); N, negative control (sdH_2_0). (B) Result of RT-PCR of samples harvested 24, 72 and 96 h post infection. M, DNA marker (MassRuler DNA Ladder Mix, Thermo Fisher, 80 to 10,000 bp).

The discovery of the viral SDH prompted us to search for other potential viral-encoded SDHA and SDHB homologs in marine metagenomes. These two subunits (SDHA and SDHB) form the catalytic core containing the redox cofactors that participate in electron transfer to ubiquinone; they are thus more conserved than SDHC and SDHD subunits. To test for the presence of this viral SDH in other viruses, we searched for *vSDHA* and *B* in marine metagenomes of the *Tara* Oceans expedition. The 50 most-similar and non-redundant SDHA and B sequences predicted from 101 *Tara* Oceans genome fragments were most likely derived from *Mimiviridae* viruses (Fig. 11). Indeed, out of 1,113 genes predicted from these 101 genome fragments, 681 were annotated at some taxonomic level, of which 449 were predicted to be cellular and 157 viral. Of the 157 viral genes, 146 and 130 had their last common ancestor in *Mimiviridae* and Mesomimivirinae, respectively. A total of 32 of the 101-genome fragments contained at least one gene predicted to be of *Mimiviridae* origin, and the larger the genome fragment, the more *Mimiviridae* genes it was found to encode (Fig. 11A). Functional analysis indicated that 12 of the 1,113 predicted genes were NCLDV hallmark genes (encoding five VLTF3s, two capsid proteins, two PCNAs, two helicases, and one PolB). The high proportion of unknown genes and genes annotated as *Mimiviridae* in the 101 *Tara* Oceans genome fragments encoding SDHA or SDHB strongly suggests that these fragments belong to *Mimiviridae* viruses. This finding demonstrates that the presence of SDH is not restricted to PkV RF01 and is arguably widespread among marine *Mimiviridae*. According to phylogenetic analyses of cellular and viral SDHA and SDHB, the viral homologs form a monophyletic group that branches deeply within eukaryotic lineages (Fig. 11B-C). Long-branch attraction bias could generate such topologies but, as explained above for the IleRS and AsnRS, it is more likely that the viral SDHA and SDHB were acquired at an early stage in the radiation of eukaryotic lineages. The transcription of *vSDHA* and its occurrence in marine environments calls for further investigation to understand the biological role and co-evolutionary significance of this viral SDH.

**FIG 11.**
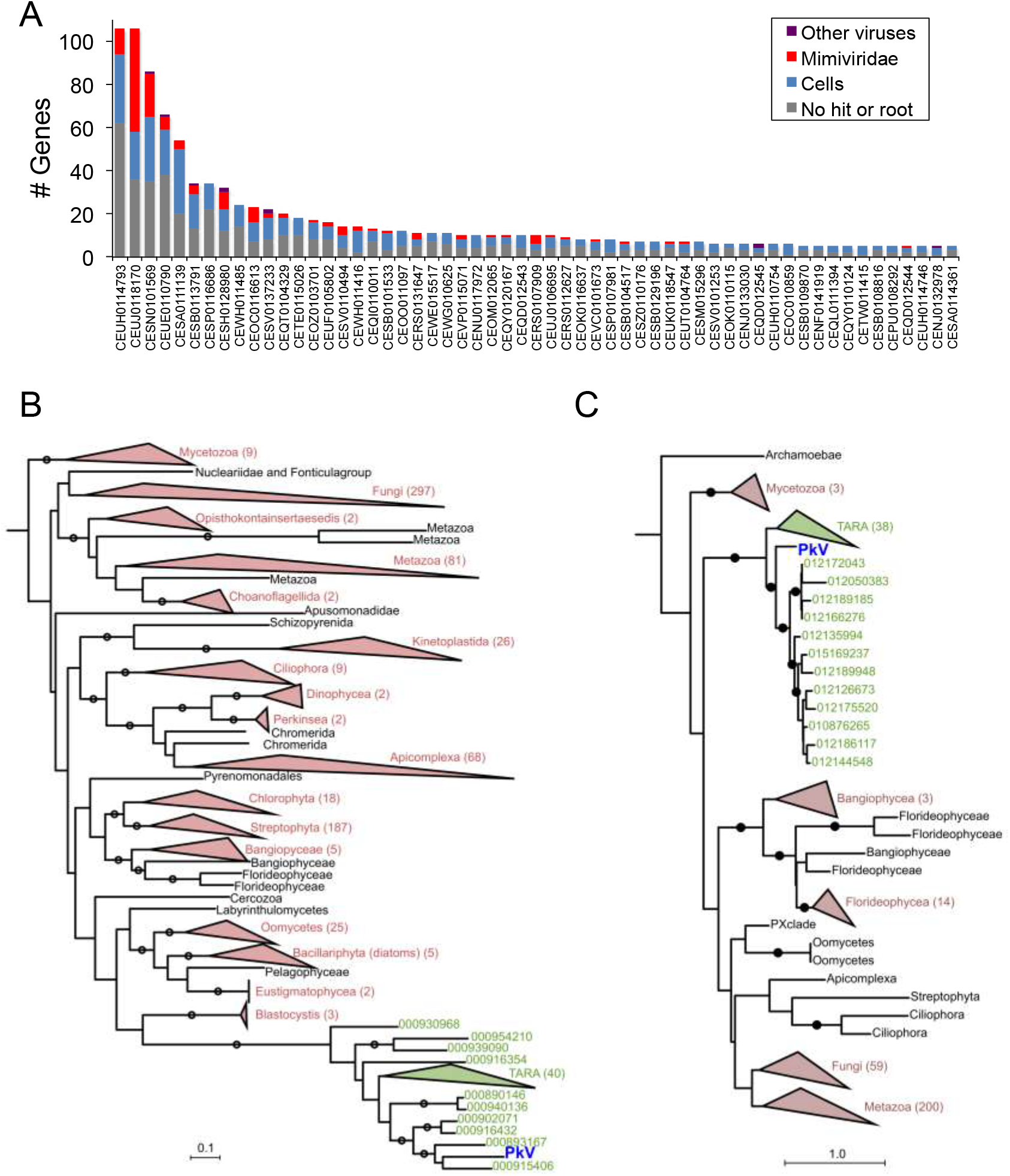
Origin of PkV RF01 SDHA and SDHB and their most similar homologs in *Tara* Oceans metagenomes. (A) Taxonomy of genes predicted in *Tara* Oceans metagenome assembled-genome fragments encoding the 50 SDHAs and SDHBs most similar to PkV RF01 genes (for genome fragments having at least five predicted genes). (B and C) Phylogenetic trees of viral and cellular SDHAs (B) and SDHBs (C). Clades in green contain PkV RF01 SDHA or SDHB and their 50 most similar hits identified in *Tara* Oceans metagenomes (predicted to be *Mimiviridae* homologs from A). Red, eukaryotic phyla; black, unclassified eukaryotes. Trees are rooted with Proteobacteria and Firmicutes homologs (not shown). Circles indicate branches with posterior probability support ≥ 50%.

Other genes related to energy production were detected in the 8,026 bp-long region. ORF 894 and ORF 896, respectively corresponding to cytochrome *c* (CytC) and cytochrome b6-f complex iron-sulfur (Cyt b6-f) subunits, showed high sequence conservation with *Chrysochromulina* sp. CCMP291 proteins (78% and 59% amino acid [aa] identities, respectively). CytC is a short protein (~100 aa) involved in the oxidative phosphorylation pathway, where it accommodates the transfer of electrons between the coenzymes Q-cytochrome *c* reductase (complex III) and cytochrome *c* oxidase (complex IV). The presence of Cyt b6-f between oxidative phosphorylation genes is puzzling because the cytochrome b6-f complex is involved in photosynthesis.

The core of the chloroplast b6f complex, however, is similar to the analogous respiratory cytochrome bc(1) complex. The other two predicted ORFs in this region are similar to ubiquinone biosynthesis protein UbiB (ORF 895) or contain a NAD-binding domain and a Fe-S cluster (ORF 897) and may thus be associated with electron transport as well. ORF 897 has two distant (25%–31% aa identity) homologs in the PkV RF01 genome (ORF 456 and ORF 625).

Some other genes were predicted to encode enzymes involved in pyruvate metabolism. ORF 79 has sequence homology with L-lactate dehydrogenases; it might thus catalyze the conversion of lactate to pyruvate, an intermediary compound serving as a starting point for several major metabolic pathways, such as glycolysis, gluconeogenesis, and the TCA cycle. ORF 727 was predicted to code for an isochorismate hydrolase that also produces pyruvate from isochorismate. ORF 24 and ORF 726 share sequence homology with phosphoenolpyruvate synthase and a partial pyruvate kinase, respectively. The former catalyzes the conversion of pyruvate to phosphoenolpyruvate (PEP), while the latter catalyzes the reverse reaction. Formation of PEP is an initial step in gluconeogenesis.

### A nearly complete viral-encoded β-oxidation pathway

In this study, 22 predicted genes were inferred to code for proteins involved in lipid synthesis or degradation, including key enzymes of the *β*-oxidation pathway (Table 2). Several genes were predicted to code for lipase-like proteins (ORFs 386, 481, 635, 653, and 690), including a triacylglycerol lipase (ORF 386) that can break down triacylglycerol into glycerol and fatty acids. Glycerol and fatty acids can be used as a starting point for ATP production—by glycolysis and *β*-oxidation, respectively. In the *β*-oxidation pathway, fatty acids are fully oxidized to produce acetyl-CoA, which can then enter the TCA cycle to yield NADH and FADH2; these latter two products can funnel through to the electron transport chain to produce ATP (Fig. 7C). Each *β*-oxidation cycle itself also produces NADH and FADH2 cofactors. We found that PkV RF01 encodes key *β*-oxidation enzymes. First, two distantly related ORFs (ORF 142 and ORF 904 sharing 22% aa identity) have sequence homology with a long-chain fatty acyl-CoA synthetase. This enzyme catalyzes the formation of fatty acyl-CoA in the cytosol. Fatty acyl-CoA can be imported to mitochondria using a (carnitine) CoA-transferase also encoded in PkV RF01 (ORF 33). Once in the mitochondrial matrix, fatty acyl-CoA serves as a substrate on which an acyl-CoA dehydrogenase (ORF 1046) oxidizes the fatty acyl-CoA and reduces a FAD cofactor to produce a FADH2 cofactor. We identified a 2,4-dienoyl-CoA reductase (ORF 30) that may facilitate the next oxidation step to produce a NADH cofactor. FADH2 and NADH molecules produced by a *β*-oxidation cycle can both be oxidized in the electron transport chain to generate ATP. The enzymes involved in the two intermediate steps following each oxidation, either an enoyl-CoA hydratase or a β-ketothiolase, were not detected in our analysis.

**TABLE 2.** Gene related to lipid metabolism.

| ORF | Annotation | KEGG orthology | Pathway |
| --- | --- | --- | --- |
| 30 | 2,4-dienoyl-CoA reductase, mitochondrial [EC:1.3.1.34] | K13236 | Beta oxidation |
| 33 | Putative CoA-transferase | NS | Beta oxidation |
| 121 | glycerophosphoryl diester phosphodiesterase | K01126 | Glycerophospholipids metabolisms |
| 138 | Fatty acid synthase (FASN) | K00665 | Fatty acid biosynthesis |
| 142 | Long-chain-fatty-acid--CoA ligase ACSBG [EC:6.2.1.3] | K15013 | Fatty acid degradation /biosynthesis / Beta Oxidation |
| 175 | Acetyl-CoA carboxylase / biotin carboxylase 1 [EC:6.4.1.2 6.3.4.14 2.1.3.15] | K11262 | Fatty acid biosynthesis |
| 236 | Glutaryl-CoA dehydrogenase [EC:1.3.8.6] | K00252 | Fatty acid degradation |
| 293 | Lysophospholipase like | NS | NS |
| 357 | Lysophospholipase like | NS | NS |
| 386 | Triacylglycerol lipase [EC:3.1.1.3] | K01046 | Glycerolipid metabolism |
| 481 | Lipase like | NS | NS |
| 635 | Lipase-like | NS | NS |
| 653 | Lipase-like | NS | NS |
| 690 | Lipase-like | NS | NS |
| 774 | Lysophospholipid Acyltransferases [EC:2.3.1.22] | K14457 | Glycerolipid metabolism |
| 694 | Lipase esterase (Carbohydrate esterase CE10) | NS | NS |
| 695 | Lipase esterase (Carbohydrate esterase CE10) | NS | NS |
| 886 | Stearoyl-CoA desaturase (Delta-9 desaturase) [EC:1.14.19.1] | K00507 | Biosynthesis of unsaturated fatty acids |
| 902 | Fatty acid synthase (FASN) | K00665 | Fatty acid biosynthesis |
| 904 | Long-chain-fatty-acid--CoA ligase ACSBG [EC:6.2.1.3] | K15013 | Fatty acid degradation /biosynthesis / Beta Oxidation |
| 1016 | Cyclopropane-fatty-acyl-phospholipid synthase [EC:2.1.1.79] | k00574 | NS |
| 1046 | Acyl-CoA dehydrogenase | K06445 | Fatty acid degradation / Beta oxidation |

Most of these genes have no homologs in reference viral genomes, and, to our knowledge, this is the first report of a virus possessing proteins directly involved in lipid-based energy production. By diverting host lipid machinery, interactions of viruses with lipids or lipid based-structures have long been known to have structural or signaling roles at different stages of the virus life cycle, such as entry, genome replication, morphogenesis, and exit (64–66). More recently, several studies on human viruses (two herpesviruses and one RNA virus) have shown that the metabolic state of an infected cell can be shifted toward energy generation to support viral replication (65). These studies have highlighted the increasing abundance—up to 48 h after HCV infection—of enzymes involved in *β*-oxidation, amino acid catabolism, and the TCA cycle (67) and an increase in cellular *β*-oxidation following the release of free fatty acids caused by Dengue virus-induced autophagy (68). Among algal viruses, EhV remodels the transcription of host lipid genes for fatty acid synthesis to support viral assembly (69) and also to generate triacylglycerols stored in the virion and available as an energy pool in later infection phases (70). Besides diverting the host metabolism, EhV encodes seven proteins involved in the sphingolipid biosynthesis pathway (71). This pathway produces a viral sphingolipid that is a central component of EhV lipid membranes and that can also act as a signaling lipid and induce programed cell death during the lytic infection phase (72). EhV also encodes a triglyceride lipase (with detectable homology to predicted PkV RF01 lipases ORF 635 and ORF653) that is highly expressed during late infection concomitantly with significant up-regulation of host *β*-oxidation genes (69). These examples and our observations of several genes involved in *β*-oxidation clearly show that viruses can introduce new metabolism-related genes, sometimes representing entire pathways, into the host, most likely to satisfy the high metabolic requirement of these giant viruses.

### High representation of glycosyltransferases

Compared with other viruses, PkV RF01 was found to encode an unusually high number of glycosyltransferases (GTs) as well as other carbohydrate-active enzymes. Automated annotation of GTs (and other carbohydrate-active enzymes) in reference viral proteomes using dbCAN2 (73) revealed that the largest number of GT domains was encoded by PkV RF01 (*n* = 48), followed by CeV (*n* = 13), *Mimivirus* members, and CroV and AaV (*n* = 8–10) (Fig. 12). We uncovered 48 GT domains encoded in 40 ORFs, 8 of which were predicted to encode more than one GT domain. These domains correspond to 16 different GT families. Most domains were inferred to be functional, as 31 out of 48 covered at least 70% of the dbCAN2 reference domain, with coverage ranging from 44% to 99%. GTs were found scattered across the genome of PkV RF01 but with some local clustering (Fig. 2A), the latter indicating possible involvement in the same pathway. GT32 was the most represented domain, with 11 proteins (as annotated by dbCAN2) and potentially three additional proteins (ORFs 40, 84, and 861). Eight proteins possessed a GT25 domain that can catalyze the transfer of various sugars onto a growing lipopolysaccharide chain during its biosynthesis. Among these eight predicted ORFs, four contained an additional non-overlapping GT domain (two GT2s, one GT6, and one GT60). Functional analyses of GTs in mimiviruses (or in related *Paramecium bursaria* Chlorella viruses) have demonstrated that some of these enzymes are functional, being able to modify viral collagen-like proteins (74) and polymerize sugars (75). Conservation between PkV RF01 GTs and functionally characterized GTs in viruses and cells is absent or extremely low, which precludes any predictions as to the specific roles of these enzymes in the PkV RF01 life cycle. Nevertheless, this putative glycosylation-conducive autonomy possibly allows the virus to infect a variety of hosts, as the virus can modify its own glycans, which are used for host recognition, independently of the host system (76). In alpha-, flavi-, and herpes-viruses, fusion is mediated by viral glycoproteins (40).

**FIG 12.**
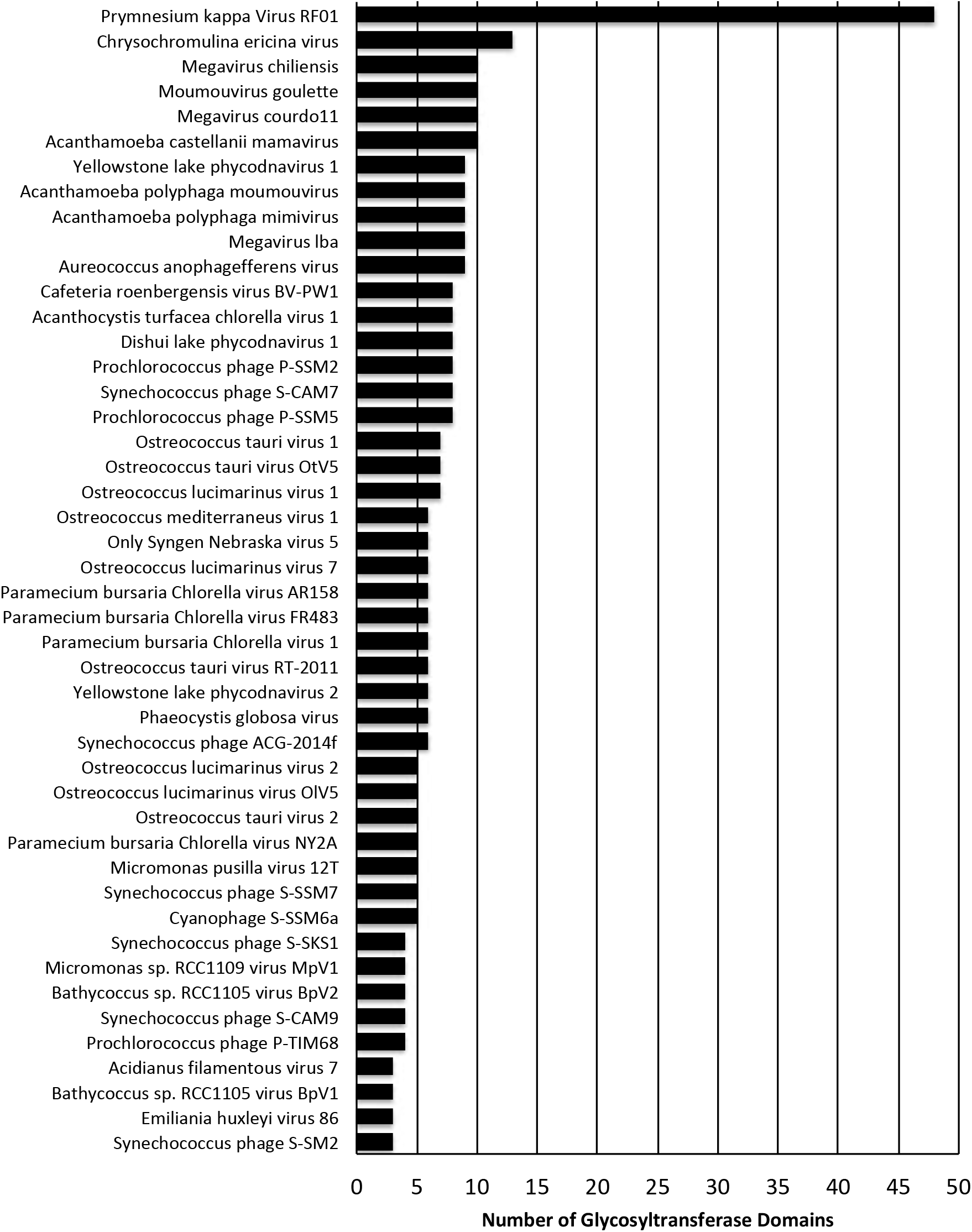
Comparative distribution of glycosyltransferase domains among viruses.

Other carbohydrate-active enzymes in the PkV RF01 genome include seven glycoside hydrolases (GHs), four carbohydrate esterases (CEs), one polysaccharide lyase (PL), one carbohydrate-binding module (CBM), and a putative sugar fermentation stimulation protein A (ORF 1003) possibly involved in maltose metabolism. These numbers are not excessively high compared with other viruses. Other detected ORFs were homologous to enzymes involved in carbohydrate transport and metabolism, notably a transketolase (ORF 528) involved in the pentose phosphate pathway in all organisms and in the Calvin cycle of photosynthetic organisms. Finally, we detected a 6-phosphofructo-2-kinase/fructose-2,6-biphosphatase 2 (ORF 539) and a mannose-1-phosphate guanylyltransferase/mannose-6-phosphate isomerase (ORF 836) respectively involved in fructose and mannose metabolism.

## Conclusions

The haptophyte virus PkV RF01 has been previously shown to have a longer replication cycle and a broader host range compared with other prymnesioviruses and most other algal viruses. Here, we revealed that PkV RF01 has atypical virion morphology and that infections yield several orders of magnitude fewer infectious particles than other tested prymnesioviruses. In-depth phylogenetic analysis using genes conserved in NCLDVs confirmed that PkV RF01 belongs to *Mimiviridae* but is deeply diverged from existing members, although closer to alga-infecting *Mimiviridae* than heterotroph-infecting ones. Unlike other alga-infecting *Mimiviridae*, however, PkV RF01 has a large genome (1.4 Mb) and contains genes coding for two aminoacyl-tRNA synthetases and the complete BER pathway. All these features are conserved in most heterotroph-infecting *Mimiviridae* and therefore must have been lost in other alga-infecting *Mimiviridae*. This outlier virus features an unprecedentedly high number of genes involved in energy metabolism and glycosylation machinery that may enable its longer replication cycle and broader host range compared with other algal viruses. These genomic and phenotypic features are suggestive of a persistent infection behavior that probably evolved in response to the host growth strategy. Because of nutrient limitations, these persistent systems of slow-growing but ubiquitous hosts with less virulent viruses may represent the most common type of virocells in oceans.

## Materials and Methods

### Culturing and infection

All algal host cultures were grown in liquid IMR/2 medium consisting of 70% aged seawater, 30% distilled water (25 PSU), and additional selenite (10 nM final concentration). The cultures were kept at 14°C and partially synchronized using a 14:10 h light: dark cycle with irradiance of 100 μmol photons m^-2^ s^-2^ supplied by white fluorescent tubes. Viruses were produced by adding freshly produced viral lysate (ca. 2 x 10^8^ VLP/mL), propagated three time on the host before added to exponentially growing host cultures (ca. 5×10^5^ cells/mL) in a ratio of 1:10 volume. Infection was followed by flow cytometry (FCM) (77, 78) for 72 h by counting viral particles and host cells, as described in (33). Burst size was calculated as the number of viral particles released from each host cell, estimated from the total number of host cells pre-infection and the total number of VLPs produced during the infection cycle (33).

### Infectious progeny

The percentage of viral infectious progeny was determined by comparing the most probable number (MPN; endpoint dilution (78)) and flow cytometric total counts of viral particles produced during infection. The number of infectious particles released in a burst was determined based on the percentage of viral infectivity produced during the infection cycle and the burst size. Infectivity was tested using *Haptolina ericin*a UiO028 as a host, and also compared with two other prymnesioviruses, HeV RF02 and PkV RF02 (33), propagated on He UiO028 and *Prymnesium kappa* RCC3423, respectively.

Briefly, 10x dilution were prepared from fresh viral lysate and added to exponentially growing host cells in 96-well microtiter plates (eight replicates for each dilution). The plates were incubated for 7 days under normal incubation conditions. Cell lysis was measured by monitoring *in situ* fluorescence on a plate reader (PerkinElmer EnSpire™ 2300 Multilabel Reader) at 460/680 nm. Numbers of infectious particles were estimated from the proportion of lysed wells using the MPN_ver4.xls excel spreadsheet from (79).

### Sensitivity to chloroform

The effect of chloroform on infectivity, used to infer the presence of a lipid membrane or lipid molecules in the capsid, was tested by adding 50% (v/v) chloroform to PkV RF01 lysate. After mixing, the chloroform phase was separated from the solution by centrifugation at 4,000 *g* for 5 min. The tubes were incubated at 37°C for 2 h with the lids open to allow evaporation of any remaining chloroform.

Triplicates of exponentially growing He UiO028 cells (1,6 x 10^5^ cells /mL) were incubated with 1:10 volumes of chloroform-treated viruses (ca. 2 x 10^8^ VLP/mL). The incubation was followed for 7 days by counting host cells by FCM (78). Host cells in chloroform-treated or untreated medium at the same ratio used with the viral lysate were used as controls. Virus propagation was confirmed in lysed cultures by FCM.

### Cryo-electron tomography

A small drop of concentrated PkV RF01 (8×109) was deposited on a glow-discharged, 200-mesh copper grid with holey carbon film (R2/1 Cu 200, Quantifoil Micro Tools GmbH, Germany). The sample was blotted with filter paper and immediately plunge frozen in liquid ethane. Grids were transferred under liquid nitrogen to a cryo-transfer tomography holder (Fishione Instruments, USA) and inserted in a 200-kV transmission electron microscope (Thermo Scientific Talos F200C) equipped with a Ceta 16M camera. Tilt series were recorded at 45,000x magnification and −7 μm defocus between −60° to 60° in 2° increments. Finally, reconstruction, segmentation, and visualization of the tomograms was performed with IMOD v4.9 software (80).

### Purification of viral particles and DNA isolation

Exponentially growing He UiO028 cultures (2 L) were infected with 20 mL of PkV RF01 and inspected visually for lysis. An uninfected culture (100 mL) was used as a control. Lysed algal cultures were checked for viruses by FCM counting. Lysed cultures were first centrifuged to remove algal debris and some bacteria (5,500 rpm for 15 min). Viruses were then pelleted by ultracentrifugation at 25,000 rpm in a Beckman Coulter Optima L90K ultracentrifuge for 2 h. The pellets were resuspended in SM buffer (0.1 M NaCl, 8 mM MgSO_4_·7H_2_0, 50 mM Tris-HCl, and 0.005% glycerin). Viral particles were further purified by Optiprep gradient centrifugation (81). Fractions were checked for viruses by FCM and for infectivity by infection of He UiO028.

Isolation of high-quality DNA for sequencing was done by following the protocol of (82) with some modifications. Viral particles were disrupted by one round of heating to 90°C for 2 min and then chilling on ice for 2 min. Disodium ethylenediaminetetraacetic acid and proteinase K at a final concentration of 20 mM and 100 μg mL^-1^, respectively, were then added before incubation of the samples for 10 min at 55°C. Sodium dodecyl sulfate at a final concentration of 0.5% (w/v) was subsequently added, and samples were incubated for an additional 1 h at 55°C.

Double-stranded DNA was then purified from the lysates using a Zymo Genomic DNA Clean & Concentrator Kit-10 (Zymo Research, Irvine, CA, USA) according to the manufacturer’s protocols. To avoid shearing DNA, gentle pipetting and mixing (accomplished by turning the tubes instead of vortexing) were performed in all steps.

### Genome assembly

Isolated DNA from PkV RF01 was subjected to Illumina TruSeq PCR-free library preparation (insert size 350 bp). The generated library was sequenced on an Illumina MiSeq instrument in paired-end mode (2 x 300 bp) to yield approximately 1.9 million reads, which corresponds to about 400x coverage. Reads were assembled into 2,498 contigs of 500 bp or more with a total assembly size of 4.75 Mb using Newbler (83). In addition, a ligation-based 1D^2^ nanopore library (LSK-308) was constructed and sequenced using an Oxford Nanopore MinION Mk1b device and a FLO-MIN107 flow cell, which resulted in 825 long reads with an N50 of 13.6 kb and a total of 9.89 Mb. To improve the assembly, short-read contigs were manually bridged with the long reads. Manual assembly using Consed (84) yielded a linear genome sequence of 1.4 Mb with inverted terminal repeats. After assembly, the consensus was polished using Nanopolish (85) and Pilon (86).

### Phylogenetic analyses

#### Five core genes, SDHA, and SDHB

The phylogenetic position of PkV RF01 was inferred from concatenated protein alignments of five core nucleocytoplasmic virus orthologous genes (NCVOGs) (87): D5-like helicase-primase (NCVOG0023), DNA polymerase elongation subunit family B (NCVOG0038), DNA or RNA helicases of superfamily II (NCVOG0076), packaging ATPase (NCVOG0249), and Poxvirus Late Transcription Factor VLTF3-like (NCVOG0262). Sequences were obtained from the NCVOG database (ftp.ncbi.nlm.nih.gov/pub/wolf/COGs/NCVOG/) (88). Additional sequences were obtained from genomes retrieved from GenBank and annotated with HMMER v3.12b using the hmmsearch (89) command with hidden Markov models available in Schulz et al. (2017) (13). Sequences from each NCVOG were aligned independently using MAFFT L-INS-i (90). The alignments were trimmed with trimAl v1.2 in *gapyout* mode (91) prior to concatenation using a custom Python script. Bayesian phylogenetic trees were inferred with PhyloBayes 1.7 (92) using the CAT model and a GTR substitution matrix. Four chains were run for 34,500–35,500 generations. The *bpcomp* command was used to check for convergence and stop when *maxdiff* = 0.3. One chain was discarded, and a consensus tree was constructed using the remaining three chains.

For phylogenetic analyses of succinate dehydrogenase subunits, top hits of PkV RF01 SDHA and SDHB were retrieved from UniProt (https://www.uniprot.org/) using online PHMMR searches (https://www.ebi.ac.uk/Tools/hmmer/search/phmmer) and also from the *Tara* Oceans project using online BLASTP searches (http://tara-oceans.mio.osupytheas.fr/ocean-gene-atlas/) (Villar et al., 2018). Alignments generated with MAFFT L-INS-i were filtered with trimAl in gapyout mode. Maximum-likelihood phylogenies were inferred with RAxML 8.2.9 (93) using the PROTCATALG model and automatic bootstrapping with the following options: ‘-N autoMRE -f a -n autoresult’. Phylogenetic trees of PkV RF01, SDHA, and SDHB were visualized using iTOL (94).

#### Rpb2, IleRS, and AsnRS

To reconstruct a phylogenetic tree based on the second largest RNA polymerase subunit, homologs were recruited by comparing Mimivirus Rpb2 against all proteins of viruses and selected organisms in the KEGG database using the GenomeNet BLASTP tool (https://www.genome.jp/). Organisms were manually selected from the KEGG list to ensure broad taxonomic coverage of the tree of life. The retrieved amino acid sequences were aligned using MAFFT-LINSI (90) and then trimmed using trimAl (91) with the following parameters: ‘-resoverlap 0.5 -seqoverlap 70 -gt 0.8 -st 0.001 -cons 50’. The tree was reconstructed using FastTree (95) as implemented in the GenomeNet TREE tool (https://www.genome.jp/tools-bin/ete). Isoleucine tRNA synthase and aspartyl tRNA synthetase viral and cellular homologs were retrieved and aligned in the same way. Trees were searched using PhyloBayes MPI (96) with the non-homogeneous CAT+GTR model (97). For each protein three chains were run until *maxdiff* parameter reach < 0.3 (0.27 for AsnRS and 0.16 for IleRS). One chain was discarded for IleRS, and a consensus tree was constructed using the remaining chains.

### Gene prediction and functional and taxonomic annotation

GeneMarkS with the option ‘virus’ (98) predicted 1,121 open reading frames (ORFs) in the fully assembled genome sequence of PkV RF01, while tRNAscan-SE (99) predicted 41 tRNAs. PkV RF01 CDS amino acid sequences were searched against Virus-Host DB (100), RefSeq (101), UniRef90 (102), and COG (61) databases using BLASTP with an *E*-value of 1 × 10^-5^ as the significant similarity threshold and against the Conserved Domain Database (103) using RPS-BLAST with an *E*-value threshold of 1 × 10^-2^. The 10 best hits for each database were compiled in a single file and manually inspected to transfer annotations of subject sequences to our query. In ambiguous cases, such as distant homologs (often seen in viral genomes) or unclear or contradictory annotations of subject sequences, the query was searched against KEGG genes (104) to allow extensive manual checking using GenomeNet tools (https://www.genome.jp/; alignment quality, length comparison to canonical genes, and links with KEGG orthology). We automatically annotated glycosyltransferases (GTs) and other carbohydrate-active enzymes (glycoside hydrolases, GHs; polysaccharide lyases, PLs; carbohydrate esterases, CEs; and auxiliary activities, AAs) in PkV RF01 and all viral genomes in Virus-Host DB (as of June 2018) using the *hmm* option of the dbCAN2 pipeline and its profile database (73). We retained hits with *E*-values < 1 × 10^-5^ and domain coverage > 35%, which corresponded to default settings.

### Taxonomic and functional analysis of vSDHA homologs in OM-RGCv1

We searched PkV RF01 SDHA and SDHB against OM-RGCv1 (105) using the Ocean Gene Atlas (106) BLAST-based tool and kept the top 50 hits with significant *E*-values for further analysis. We then collected genome fragments (contigs) encoding these 50 SDHAs and 50 SDHBs by searching via BLASTN for identical hits over full *SDHA* or *SDHB* lengths against *Tara* ocean assemblies (downloaded from EBI) used to construct OM-RGCv1. We predicted ORFs in these genome fragments using GeneMarkS. The resulting 1,113 amino acid sequences were functionally annotated by searching against Pfam protein families (107) using profile HMM scan (108) and also taxonomically using a last common ancestor strategy as in (109); in brief, protein sequences were searched against a database composed of UniRef cells, MMETSP (110) and Virus-Host DB (100) data using DIAMOND (111). Selected hits were then used to derive the last common ancestor of the query using a NCBI taxonomic tree re-wired to reflect the taxonomy of NCLDVs.

### PCR and RT-PCR optimization

We designed specific primers (Table 3) targeting a 256-bp region of the *mcp* gene to use both as an internal control in the RT-PCR and to confirm that our protocols were optimized. For each PCR, a negative control (sterile distilled H_2_O) was included. PCR amplifications were carried out in 50-μL total volumes containing 1 μL of template using a DNA HotStarTaq Master Mix kit (Qiagen). The cycling protocol was as follows: 15 min at 95°C, followed by 35 cycles of 30 s at 94°C, 30 s at 59°C, and 30 s at 72°C, with a final extension of 12 min at 72°C.

**TABLE 3.**
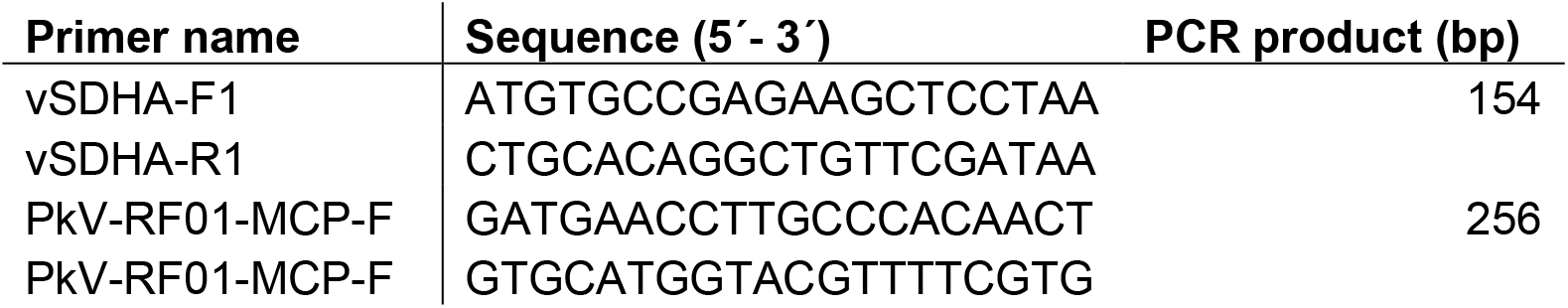
Forward and reverse PCR primers for amplification of vSDHA and MCP

RT-PCRs were performed using the SuperScript III One-Step RT-PCR with Platinum *Taq* DNA Polymerase system (Thermo Fisher). Cycling conditions were as follows: 16 min at 55°C and 2 min at 94°C, followed by 40 cycles of 15 s at 94°C, 30 s at 49°C, and 30 s at 68°C, and a final extension of 5 min at 68°C.

All PCR products were checked for the correct size on a 1.5% agarose gel stained with GelRed (Biotium). PCR products were further checked by sequencing using BigDye v3.1 (Thermo Fisher) for cycle sequencing (Sekvenseringslaboratoriet, UiB, Norway).

### PCR amplification and RT-PCR analysis of *vSDHA*

To investigate whether the *vSDHA* gene is transcribed during infection, an infected culture of He_UiO028 plus PkV RF01 as well as an uninfected He UiO028 culture (control) were set up as described above. Samples were collected at 24, 72, and 96 h post infection from both cultures. RNA was extracted using an RNeasy Plus Universal Mini kit (Qiagen), with gDNA removed in an extra step using a TURBO DNA-free kit (Ambion).

Specific primers were designed to target a 150-bp region of the *vSDHA* gene (Table 3). For each PCR, two negative controls (sterile distilled H_2_O and extracted DNA from He028) were included. As positive controls for the transcription, we used primers targeting the *mcp* gene (see above). As a positive PCR control, we used genomic PkV RF01 DNA. PCR amplifications were conducted in 50-μL total volumes containing 1 μL of template DNA using an ExTaq kit (Takara). The cycling protocol was as follows: 5 min at 94°C, followed by 35 cycles of 30 s at 94°C, 30 s at 59°C, and 30 s at 72°C, with a final extension of 12 min extension at 72°C.

RT-PCRs were performed using a SuperScript III One-Step RT-PCR with Platinum Taq DNA Polymerase system (Thermo Fisher). Cycling conditions were as follows: 16 min at 55°C and 2 min at 94°C, followed by 40 cycles of 15 s at 94°C, 30 s at 49°C, and 30 s at 68°C, with a final extension of 5 min at 68°C. PCR products were checked as described above.

## Data availability

Raw sequence reads and PkV RF01 genome sequence were deposited at the European Bioinformatics Institute (EMBL-EBI) (https://www.ebi.ac.uk) under project name PRJEB37450. The complete video records of a cryo-electron tomogram of a PkV RF01 virion and sequence data as well as curated gene annotation table as reported in this study are available at https://github.com/RomainBlancMathieu/PkV-RF01.

## Acknowledgements

The recording of tilt series was performed with the help of Sebastian Schultz at the Unit of Cellular Electron Microscopy, the Norwegian Radium Hospital. Initial sequencing (MiSeq and Pacbio) of PkV RF01 total DNA was performed at the Norwegian Sequencing Center (https://www.sequencing.uio.no/). We thank Hilde M. K. Stabell and Solveig Siqveland, Department of Biological Sciences, University of Bergen, Norway, for technical assistance with molecular biology experiments as well as Christian Rückert, Bielefeld University, for support in manual finishing of genome assembly and Minyue Fan, Kyoto University, for assistance in genes analysis. This work was supported by the Research Council of Norway project entitled “Uncovering the key players for regulation of phytoplankton function and structure: lesson to be learned from algal virus-haptophyte coexistence” (VirVar, project number 294364 to RAS). Additional funding was provided by the European Union Horizons 2020 research and innovation program, grant agreement no. 685778 (“Virus-X”) to RAS and DB. This work was also supported by the Future Development Funding Program of the Kyoto University Research Coordination Alliance. HO was supported by JSPS/KAKENHI (No. 18H02279), and Scientific Research on Innovative Areas from the Ministry of Education, Culture, Science, Sports and Technology (MEXT) of Japan (Nos. 16H06429, 16K21723, 16H06437). The Super Computer System, Institute for Chemical Research, Kyoto University, provided computational time. We thank Barbara Goodson, from Edanz Group (www.edanzediting.com/ac), for editing the English text of a draft of this manuscript.

